# Longitudinal fMRI reveals altered developmental trajectories of functional network organisation associated with adolescent cannabis use

**DOI:** 10.64898/2026.10.20.760368

**Authors:** Elizabeth J. Davies, Natalie Ertl, Tom P. Freeman, Anya Borissova, Rachel Lees Thorne, Shelan Ofori, Kat Petrilli, Clare Mokrysz, H. Valerie Curran, Will Lawn, Matthew B. Wall

## Abstract

Adolescence is a critical period of brain development during which the endocannabinoid system continues to mature. These developmental changes may contribute to age-related differences in the association between cannabis use and large-scale functional network organisation, yet this association remains incompletely characterised. We hypothesised that regular cannabis use would be associated with differential one-year changes in resting-state functional connectivity between adolescents and adults. Resting-state fMRI data at two time points, one year apart, were analysed from 66 participants (31 aged 16-17 years; 35 aged 26-29 years) including 29 with regular cannabis use and 37 age-matched cannabis-naïve controls. Seed-to-voxel analyses examined longitudinal changes in functional connectivity across age and cannabis-use groups. Year change, age × user-group interactions were identified in salience and default mode network connectivity, including divergent changes in anticorrelations with medial prefrontal, thalamic, cerebellar and juxtapositional regions. Directionality of effects differed by network and region. Additional main effects of user group were observed for executive control and associative striatal connectivity, with the cannabis-use group showing decreased positive connectivity and reduced anticorrelation, respectively. Changes in functional connectivity were developmentally dissociable and predominantly observed in regions contributing to attentional and transmodal processing. These findings suggest that cannabis may be associated with alterations in the maturational refinement of higher-order neural systems, potentially reflecting age-related differences in neurobiological sensitivity to cannabis exposure. Considering developmental stage may therefore be important for characterising cannabis-related neurobiological differences and may have implications for understanding age-dependent vulnerability to cannabis-related psychiatric outcomes.

## Introduction

Western shifts in drug policy reflect increasingly liberal public discourse (Freeman et al., 2026; London Drugs Commission, 2025) which may contribute to increased perceived social acceptability, lower perceived risk and influence patterns of uptake among younger populations (Jessor, 1991; Parker, 2005). International cannabis markets are currently dominated by high-potency strains characterised by elevated Δ^9^-tetrahydrocannabinol (THC) and low cannabidiol (CBD) levels (Freeman et al., 2021). High-potency strains increase the likelihood of problematic use and treatment need (Forti et al., 2019; Petrilli et al., 2022). Importantly, adolescence is itself a risk factor for current and future problematic cannabis use (Le Strat et al., 2015; Skumlien et al., 2025; Winters & Lee, 2008). This confluence of factors highlights an urgent need for greater understanding of the effects of chronic cannabis use on developmental brain function.

Adolescence represents a critical neurodevelopmental window characterised by asynchronous maturation of large-scale brain networks; for instance, subcortical reward systems mature earlier than prefrontal regions that are known to support cognitive control (Bossong & Niesink, 2010; Gogtay et al., 2004). These neural networks facilitate different processes and develop non-linearly, fostering changes to connectivity characteristics both within and between networks. Specifically, regions of cognitive control in the prefrontal cortex (PFC) become more specialised and segregated, whereas the salience network becomes more integrative (Gracia-Tabuenca et al., 2021; Lopez et al., 2020; Marek et al., 2015). Furthermore, normative maturation of such networks underlies executive functioning improvements observed with age (Baum et al., 2017; Marek et al., 2015; Rubia, 2013).

Similarly, the endocannabinoid system (ECS), a widely distributed neuromodulatory system, undergoes dynamic developmental changes during this period. Among its various roles, the ECS regulates synaptic transmission and plasticity, with cannabinoid receptor type 1 (CB1) mediating predominantly presynaptic control of neurotransmitter release (Hashimotodani et al., 2007). CB1 receptors are highly expressed across prefrontal, striatal, and limbic regions (Glass et al., 1997). In animal models, CB1 receptor expression peaks around the onset of adolescence before declining towards adulthood, while endogenous cannabinoid signalling shows regionally and temporally specific changes across this period (Heng et al., 2011; Meyer et al., 2018). These developmental changes in ECS signalling occur alongside neural circuit maturation, including changes in synaptic plasticity and excitatory–inhibitory regulation, while experimental disruption of cannabinoid signalling during development can alter subsequent neural and behavioural outcomes (Mulder et al., 2008; Wu et al., 2010).

The primary psychoactive component of cannabis (THC) acts primarily through partial agonism at CB1 receptors. Chronic THC exposure can induce CB1 receptor downregulation and alter neuromodulatory signalling (Cohen et al., 2019; Hirvonen et al., 2012; Sim-Selley et al., 2006). In adults, these effects appear reversible with abstinence; however, cannabis exposure during adolescence may disrupt normative patterns of neurodevelopment and consequently alter functional network stabilisation (Bossong & Niesink, 2010; Mato et al., 2004; Rubino et al., 2015).

Resting-state functional MRI (rs-fMRI) measures spontaneous low-frequency blood-oxygenation-level-dependent (BOLD) fluctuations, from which intrinsic functional connectivity (FC) and large-scale brain network organisation can be inferred (Biswal & Uddin, 2025). As a task-free imaging modality, rs-fMRI, can predict individual differences in neural activity (Cole et al., 2016; Tavor et al., 2016), and identify distributed cortical and subcortical networks (Biswal & Uddin, 2025; Greicius et al., 2003). Consequently, rs-fMRI is well suited for studying adolescent neurodevelopment and its modulation by environmental factors such as cannabis use. Using rs-fMRI, Marek et al. (2015) demonstrated that connectivity within and between large-scale networks changes non-linearly across adolescence. For example, normative maturation is characterised by increasing segregation of the default mode network (DMN), which supports internally directed cognition (Greicius et al., 2003; Smallwood et al., 2021), from task-positive networks including the executive control (ECN) and salience (SN) networks, resulting in stronger anticorrelations (negative functional coupling) between these systems (Marsh et al., 2006; Rubia, 2013; Uddin et al., 2009). The SN detects salient stimuli and is thought to facilitate switching between internally oriented (DMN) and externally directed (ECN) processing (Goulden et al., 2014; Menon, 2011; Seeley et al., 2007). Beyond cortical networks, hippocampal and striatal systems contribute to large-scale organisation, with striatal subdivisions forming partially segregated corticostriatal loops supporting reward-motivated (limbic striatum), goal-directed (associative striatum), and motor control processes (sensorimotor; Di Martino et al., 2008; Joel & Weiner, 2000).

Reported alterations as a result of cannabis use in rs-fMRI FC studies most frequently are observed across transmodal regions implicating ECN, DMN, SN, and limbic systems (Batalla et al., 2013; Ertl et al., 2023; Thomson et al., 2022; Wetherill et al., 2015; Zhou et al., 2018). However, findings vary considerably across studies, with inconsistent effects limiting convergence across meta-analyses (Thomson et al., 2022). This heterogeneity likely reflects methodological differences, including variation in developmental stage, use frequency, onset and severity, abstinence duration, co-occurring substance use, and statistical power. Pre-scan abstinence periods, for example, can modulate neuropharmacological outcomes (Hirvonen et al., 2012). Conflation of developmental stages, e.g. combining adolescent and adult populations, may obscure potential effects by failing to account for differences in neural network maturation. Among studies that better account for age, alterations are more consistently observed within higher-order attentional and control networks (Thomson et al., 2022). For example, reduced salience–parietal connectivity has been reported in adolescent and young adult users (Harris et al., 2022), while ECN-specific alterations with limited involvement of other networks have also been observed (Ertl et al., 2023b). In adults, cannabis use has been associated with altered DMN–SN coupling, and changes to within– and between-network integration including anticorrelational relationships (Pujol et al., 2014; Thomson et al., 2022). Other reports have shown differential activation of prefrontal attentional and cerebellar networks in chronic cannabis users, which correlated with age of first use (Chang et al., 2006). Moreover, earlier onset of cannabis use is an established risk factor for subsequent psychotic symptoms, even in non-clinical samples, underscoring this developmental window of potential vulnerability (Arseneault et al., 2004; Stefanis et al., 2004).

Despite this growing evidence base, no study has directly compared longitudinal developmental trajectories of resting-state connectivity in matched groups of adolescents and adults who do and do not use cannabis. This gap is particularly important given that neurodevelopment continues into the mid-twenties (Arain et al., 2013) and that endocannabinoid system activity peaks phenotypically in adolescence; it is therefore plausible that the impact of chronic THC exposure differs between age groups.

The present study addresses these gaps using a longitudinal design examining resting-state network connectivity in adolescent (16–17 years) and adult (26–29 years) weekly cannabis users and age-matched controls. We hypothesised that regular cannabis use would be associated with longitudinal changes in resting-state network connectivity, with an age × user group interaction such that adolescent users would show distinct trajectories relative to both adult users and adolescent controls. It was further hypothesised that connectivity change would be associated with cumulative cannabis exposure across the study period.

## Methods

### Ethics

The study was approved by the University College London Research Ethics Committee (Project ID: 5929/003) and conducted in accordance with the Declaration of Helsinki. All participants provided written informed consent prior to participation.

### Participants

The current analyses derive from the longitudinal arm of the CannTeen study, a prospective cohort examining neuropsychiatric outcomes in cannabis users across adolescence and early adulthood (Lawn et al., 2019; Lawn, Fernandez-Vinson, et al., 2022; Lawn, Mokrysz, et al., 2022). This analysis focused on rs-FC, contrasting consistent cannabis users and non-using controls across two developmental stages. Participants completed two MRI visits: baseline (timepoint; TP1) and one-year follow-up (TP2; mean interval = 379 days). Behavioural assessments were conducted at three-month intervals across the same period, for up to five sessions including baseline. Participants were recruited as described in (Lawn, et al., 2022).

Adolescents were aged 16–17 years and adults 26–29 years at enrolment. The ‘cannabis user’ group required use of cannabis at least once per week, averaged over the preceding three months, at each attended study session. Adult users were excluded at baseline if they reported using cannabis once or more per week prior to age 18. Controls were eligible if they had 1–10 occasions of cannabis exposure, or no cannabis use and at least one day of lifetime cigarette use, to approximate users in factors associated with opportunity for substance exposure. Participants were excluded if they were: taking psychotropic medication, receiving treatment for a mental health disorder, had a history of psychotic disorder, or used any single illicit drug other than cannabis more than two days per month in the study period.

Prior to each MRI visit, participants abstained from cannabis and alcohol for at least 12 hours and other illicit substances for 48 hours, verified via salivary toxicology on the scan day.

Of 139 participants who completed the baseline MRI visit, exclusions due to attrition, excessive head motion, incomplete imaging data, and failures to maintain the predefined longitudinal grouping criteria (cannabis usage) resulted in a final analytic sample of 66 participants (consistent cannabis users: n = 29; consistent controls: n = 37; adolescents: n = 31; adults: n = 35). A participant flow diagram is shown in Figure 1 with a breakdown of age and sex by user group.

**Figure 1.**
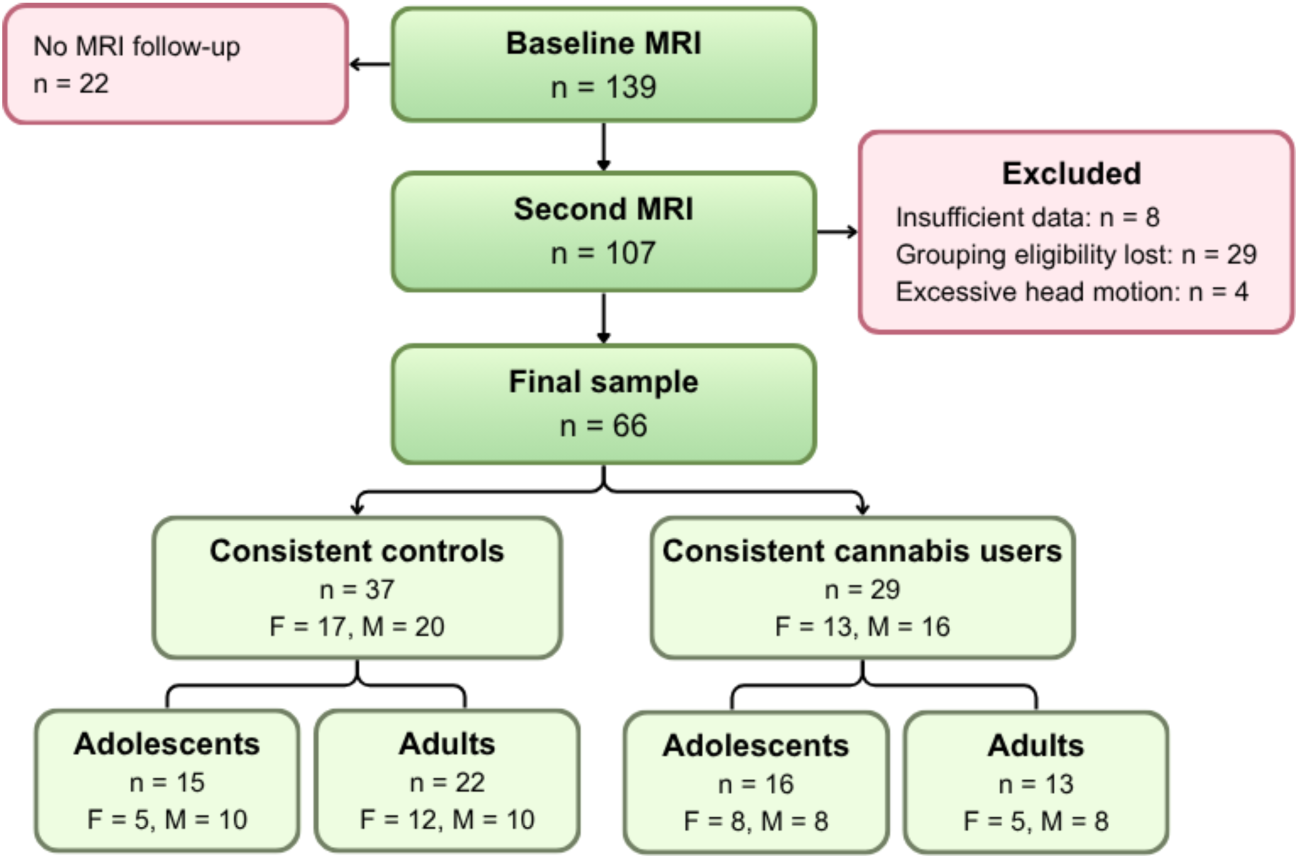
Participant flow diagram starting with baseline MRI attending sample who met initial inclusion criteria (see Lawn et al., 2019). Chart shows stages of attrition through the study reaching the final analytic sample split by user group, age group and sex.

### MRI acquisition

MRI data were collected at Perceptive Inc. (formerly Invicro; Hammersmith Hospital, London, UK), using a Siemens Magnetom Verio 3T scanner (Erlangen, Germany) and a 32-channel phased array head coil. T1-weighted structural images were acquired using a magnetisation-prepared rapid gradient-echo sequence at 1 mm resolution. rs-fMRI data were acquired following structural imaging. T2*-weighted functional images were acquired using a multiband gradient-echo, echo-planar imaging sequence (3 mm resolution; 8 minutes in duration). See Supplementary Material S1 for structural and functional scan parameters.

### MRI preprocessing

T1-weighted images were preprocessed in FSL (Smith et al., 2004) using a standard workflow (Ertl et al., 2023b). Steps included tissue segmentation to produce partial volume maps and voxel-wise field maps were generated from complex-valued MRI data. Full details of preprocessing procedures are in the Supplementary Material S2 for structural and functional data. Using FEAT, functional images were preprocessed similarly to conventions (Ertl et al., 2023).

White matter (WM) and cerebro-spinal fluid (CSF) masks were generated by transforming segmented T1 partial volume maps to each participant’s functional space, thresholding at 0.5, and then binarizing. The masks were used to extract the mean WM and CSF time series, for inclusion as nuisance regressors.

Seed regions for 7 networks were defined in MNI152 space using a mixed strategy, including meta-analytic maps from Neurosynth, the Harvard–Oxford atlas, and definitions from literature (Martinez et al., 2003; Tziortzi et al., 2014). Table 1 contains the networks of interest and corresponding seeds with centre-of-gravity coordinates in MNI space. Networks and corresponding *a priori* seed regions were the same as used in a previous baseline cross-sectional analysis of these data (Ertl et al. 2023), these seeds are visualised in Supplementary Figure S1. Standard-space binary masks were transformed to each subject’s functional space, thresholded at 0.5, binarized, and used to extract the mean time series for each network’s seed.

**Table 1.**
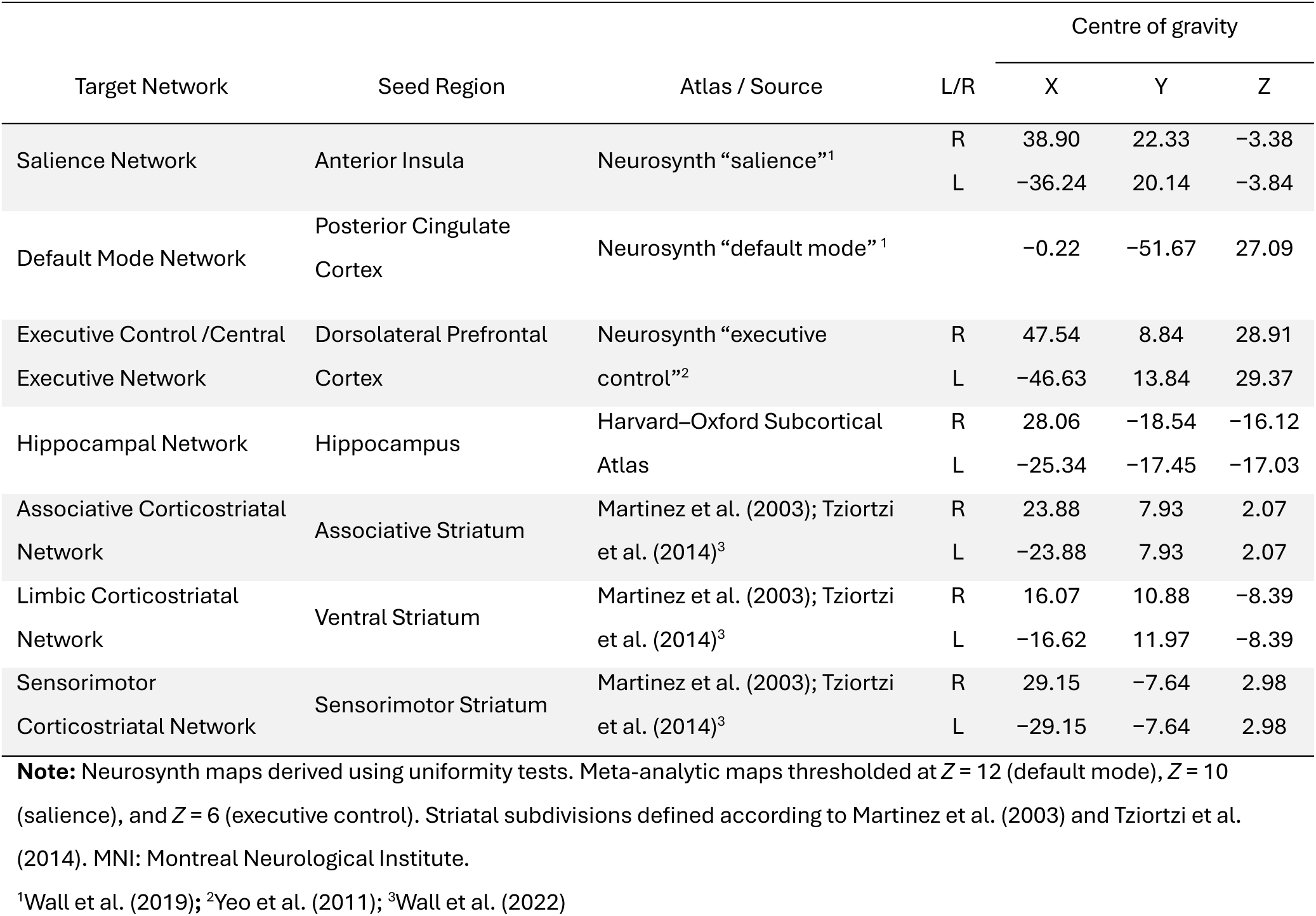
Approximate bilateral centre of gravity coordinates in MNI152 standard space of the seven network seeds used for analysis of resting state networks.

### Resting state fMRI data analysis

First-level connectivity maps were generated separately for each participant, visit, and network using a general linear model in FEAT (Woolrich et al., 2001). For each visit, voxelwise BOLD time series were modelled against the corresponding seed-region time series and prewhitened to account for temporal autocorrelation. WM and CSF regressors were included for denoising, alongside an extended 24-parameter motion model derived from MCFLIRT estimates (6 rigid-body motion parameters, their temporal derivatives, and squared terms) to minimise motion-related confounds.

To model within-subject longitudinal change in FC, first-level results from each participant’s two visits were entered into a fixed-effects mid-level analysis (FEAT). For each seed, voxelwise contrasts representing change between visits (TP2 > TP1) were generated and used in group-level analyses.

Group-level analyses modelled the main effects of user group (cannabis users vs controls), age group (adolescents vs adults), and their interaction on the mid-level change (TP2 > TP1) maps. Interaction effects were assessed using *F*-contrasts. Statistical maps were thresholded at a cluster-forming threshold of *Z* > 2.3 and cluster level-correction for multiple comparisons of *p* < 0.05.

For further analyses and visualisations, binary masks were created from significant clusters. These masks were applied at the subject level to extract mean voxelwise connectivity estimates within each cluster at TP1 and TP2, enabling characterisation of the directionality of effects (positive correlation or anticorrelation with the seed region). In addition, cluster-based connectivity change values were extracted from mid-level maps (TP2 > TP1) to visualise directionality as bar plots (generated in Python using Matplotlib; Hunter, 2007).

Exploratory linear regressions were conducted within the user group (n = 29) to probe significant main and interaction effects, relating mean connectivity estimates extracted from significant clusters showing longitudinal change to cannabis exposure metrics. Cannabis exposure was quantified in two ways: (i) a time-weighted frequency (average number of use days per week) across four 12-week intervals during which participants attended assessments, and (ii) this measure multiplied by estimated grams per week.

Behavioural and demographic statistics were assessed using *t*-tests, Mann–Whitney U tests, ANOVAs, and chi-squared tests as appropriate, using Jamovi (The jamovi project, 2023).

## Results

### Participant Characteristics

Participant demographics are in Table 2 below. There were no significant differences among groups in sex, ethnicity, or education (results of formal statistical testing are in Supplementary Material S3). Cannabis users were more likely to smoke tobacco than controls with a 93% vs 38% prevalence respectively, *χ*²(66)= 26.70, *p* < 0.001.

**Table 2.**
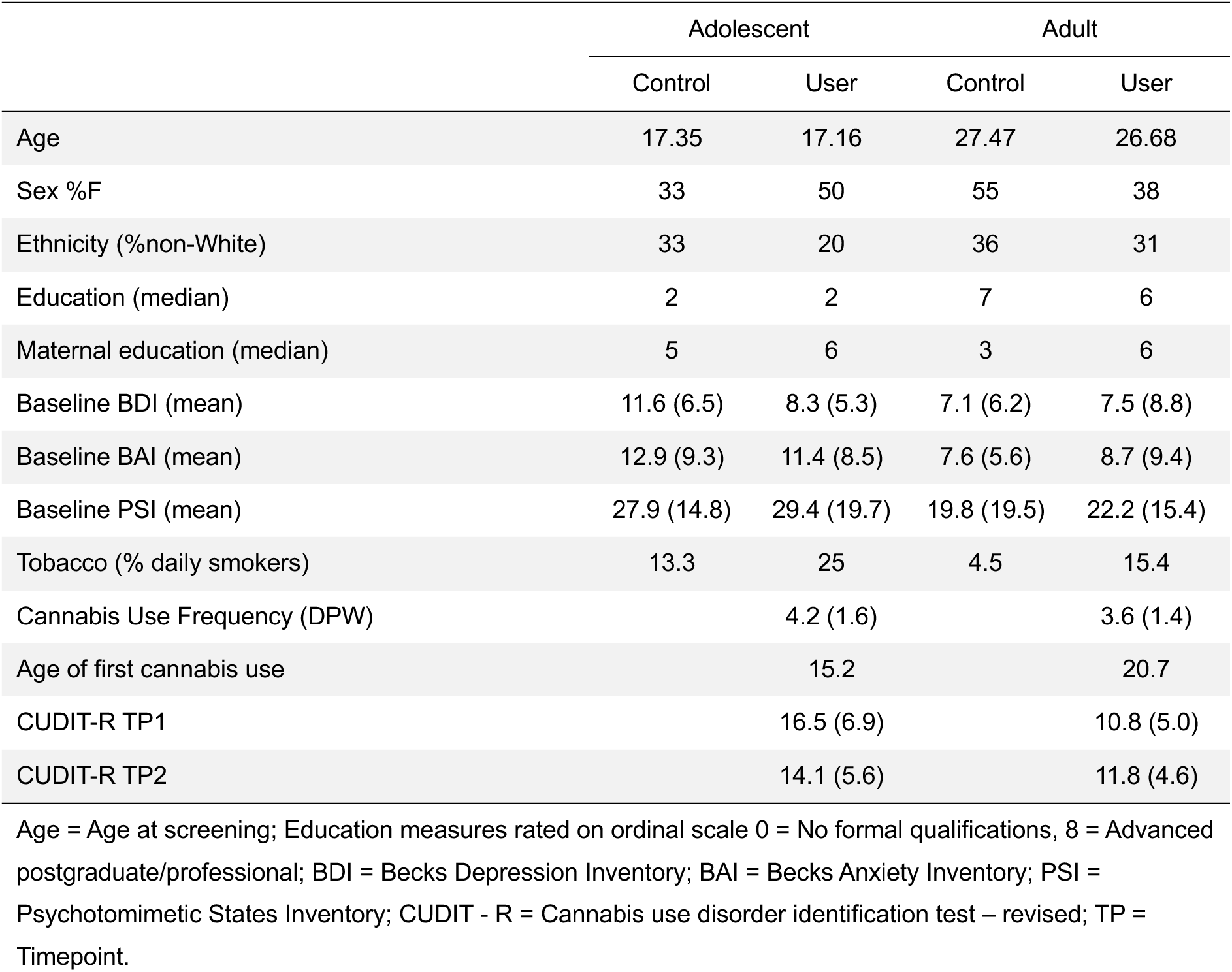
Participant demographics of the final analytic sample (n = 66) organised by subgroup.

|  | Adolescent |  | Adult |  |
| --- | --- | --- | --- | --- |
|  | Control | User | Control | User |
| Age | 17.35 | 17.16 | 27.47 | 26.68 |
| Sex %F | 33 | 50 | 55 | 38 |
| Ethnicity (%non-White) | 33 | 20 | 36 | 31 |
| Education (median) | 2 | 2 | 7 | 6 |
| Maternal education (median) | 5 | 6 | 3 | 6 |
| Baseline BDI (mean) | 11.6 (6.5) | 8.3 (5.3) | 7.1 (6.2) | 7.5 (8.8) |
| Baseline BAI (mean) | 12.9 (9.3) | 11.4 (8.5) | 7.6 (5.6) | 8.7 (9.4) |
| Baseline PSI (mean) | 27.9 (14.8) | 29.4 (19.7) | 19.8 (19.5) | 22.2 (15.4) |
| Tobacco (% daily smokers) | 13.3 | 25 | 4.5 | 15.4 |
| Cannabis Use Frequency (DPW) |  | 4.2 (1.6) |  | 3.6 (1.4) |
| Age of first cannabis use |  | 15.2 |  | 20.7 |
| CUDIT-R TP1 |  | 16.5 (6.9) |  | 10.8 (5.0) |
| CUDIT-R TP2 |  | 14.1 (5.6) |  | 11.8 (4.6) |

Age = Age at screening; Education measures rated on ordinal scale 0 = No formal qualifications, 8 = Advanced postgraduate/professional; BDI = Becks Depression Inventory; BAI = Becks Anxiety Inventory; PSI = Psychotomimetic States Inventory; CUDIT - R = Cannabis use disorder identification test – revised; TP = Timepoint.

Across visits, cannabis users consumed cannabis a mean of 3.9 ± 1.6 days/week (equating to 2.3 ± 2.5 g/week). Cannabis use in days per week was the same between age groups (*t*(27) = 1.20, *p* = 0.240). Adolescents reported higher baseline cannabis use disorder identification test – revised (CUDIT-R) scores than adults *t*(27)= −2.48, *p* = 0.020. A repeated-measures ANOVA examining CUDIT-R scores across TP1 and TP2 showed no significant main effect of age group, *F*(1,25) = 3.79, *p* = 0.063 or time, *F*(1,25) = 0.278, *p* = 0.603, and no significant age group × time interaction, *F*(1,25) = 1.169, *p* = 0.290. Results from formal statistical testing of substance use (alcohol and tobacco) and other psychological measures are in Supplementary Material S3.

Data quality control measures are in Supplementary Material S4.

### Seed-to-voxel analyses

Whole-brain seed-to-voxel analyses were performed on TP1–TP2 change maps (n=66) and parameter estimates were extracted from significant clusters. Directionality was visualised according to age group (adolescent vs. adult) and cannabis user group (user vs. control).

All significant effects are reported in Table 3 for positive FC and Table 4 for anticorrelations. A composite table of results for all network analyses is in Supplementary Table S10. No main or interaction effects were identified in the limbic or sensorimotor analyses.

**Table 3.** Significant clusters from seed-to-voxel resting-state functional connectivity analyses (cluster-corrected *p*_(FWE)_ < .05) of regions with a positive functional connectivity to the seed region.

| Network | Cluster Size<br>(voxels) | L/R | Region | Direction<br>of change | Peak MNI |  |  | Peak Z | Cluster<br>*p*(FWE) |
| --- | --- | --- | --- | --- | --- | --- | --- | --- | --- |
|  |  |  |  |  | x | y | z |  |  |
| Main Effect of Age |  |  |  |  |  |  |  |  |  |
| Associative<br>striatal | 1730 | R | Dorsolateral prefrontal cortex extending lateral precentral gyrus / premotor cortex | A ↑, T ↓ | 60 | 10 | 46 | 3.78 | < 0.001 |
|  | 493 | L | Precentral gyrus extending inferior parietal lobule and supramarginal gyrus | A ↑, T ↓ | −40 | −10 | 34 | 3.89 | 0.016 |
|  | 478 | L | Medial premotor cortex | A ↑, T ↓ | −6 | −8 | 58 | 3.86 | 0.019 |
| Hippocampal | 857 | R | Temporo-parietal junction encompassing angular gyrus (PGp) and lateral occipital cortex | A ↑, T ↓ | 50 | −64 | 26 | 4.18 | < 0.001 |
|  | 419 | L | Ventrolateral prefrontal cortex and inferior frontal gyrus extending lateral orbitofrontal cortex | A ↑, T ↓ | −44 | 18 | 10 | 3.36 | 0.048 |
| Main Effect of Cannabis Use |  |  |  |  |  |  |  |  |  |
| ECN | 714 | L | Superior parietal lobule and medial precuneus extending postcentral gyrus | C ↑, U ↓ | −8 | −42 | 56 | 3.85 | 0.004 |
|  | 500 | R | Superior parietal lobule and medial precuneus extending medial postcentral gyrus | C ↑, U ↓ | 16 | −54 | 52 | 3.69 | 0.031 |

Note: Results reflect longitudinal change over a one-year follow-up interval. MNI coordinates reflect cluster peak. Direction of change: A = Adults, T = Teenagers(/Adolescents), C = Controls, U = Cannabis users; ↑ = increasing connectivity, ↓ = decreasing connectivity over the year. ECN = executive control network; SN = salience network; DMN = default mode network; mPFC = medial prefrontal cortex; FWE = family-wise error.

**Table 4.** Significant clusters from seed-to-voxel resting-state functional connectivity analyses (cluster-corrected *p*_(FWE)_ < .05) of regions anticorrelated (negative functional connectivity) to the seed region.

| Network | Cluster Size<br>(voxels) | L/R | Region | Direction of<br>change* | Peak MNI |  |  | Peak Z | Cluster<br>*p*(FWE) |
| --- | --- | --- | --- | --- | --- | --- | --- | --- | --- |
|  |  |  |  |  | x | y | z |  |  |
| Main Effect of Age |  |  |  |  |  |  |  |  |  |
| Hippocampal | 460 | L | Lateral anterior supramarginal gyrus | A ↑, T ↓ | −64 | −32 | 46 | 3.81 | 0.030 |
| Main Effect of Cannabis Use |  |  |  |  |  |  |  |  |  |
| Associative<br>striatal | 1062 | R | Parieto-occipital sulcus extending cuneal<br>and calcarine regions | C ↓, U ↑ | 18 | −68 | 16 | 3.85 | < 0.001 |
|  | 418 | L | Middle temporal gyrus (temporooccipital) /<br>inferior lateral occipital cortex | C ↓, U ↑ | −58 | −58 | 8 | 3.59 | 0.039 |
| Age × Cannabis Use Group Interaction |  |  |  |  |  |  |  |  |  |
| SN | 1387 | L | Dorsomedial prefrontal cortex | AC+TU ↑<br>AU+TC ↓ | −12 | 62 | 34 | 4.13 | < 0.001 |
|  | 473 | R | Dorsomedial thalamus | AU+TC ↑<br>TU+AC ↓ | 6 | −24 | 12 | 4.08 | 0.024 |
|  | 424 | R | Ventromedial prefrontal cortex | AC+TU ↑<br>AU+TC ↓ | −6 | 42 | 58 | 3.32 | 0.042 |
| DMN | 509 |  | Midline anterior cerebellum lobules V-VI | AU+TC ↑<br>TU+AC ↓ | −4 | −66 | −14 | 3.94 | 0.029 |
|  | 458 | R | Pre-supplementary motor area | AC+TU ↑<br>AU+TC ↓ | 6 | 8 | 52 | 3.74 | 0.049 |

Note: Results reflect longitudinal change over a one-year follow-up interval. MNI coordinates reflect cluster peak. Direction of change: A = Adults, T = Teenagers(/Adolescents), C = Controls, U = Cannabis users; \* ↑ = increasing connectivity / reduced anticorrelation, ↓ = decreasing connectivity / strengthened anticorrelation over the year. ECN = executive control network; SN = salience network; DMN = default mode network; mPFC = medial prefrontal cortex; FWE = family-wise error.

### Age group main effects

A significant main effect of age group on FC changes (TP2 > TP1) was observed in the associative striatal and hippocampal networks, with three significant clusters identified in each. Across these effects, FC values increased in adults and decreased in adolescents over the year.

In the hippocampal seed-based analysis, significant clusters were identified in the temporo-parietal junction extending into the angular gyrus and the left ventrolateral PFC extending to the lateral orbitofrontal cortex (Table 3). An anticorrelated cluster was observed in the left supramarginal gyrus, here, an increase or decrease in FC reflected a shift toward weaker anticorrelation in adults and stronger anticorrelation in adolescents, respectively (Table 4). Figure 2 illustrates the significant clusters and subgroup FC trajectories over the year as bar charts. Directionality of effect plots for all other significant clusters are in Supplementary Figures S2–5.

**Figure 2.**
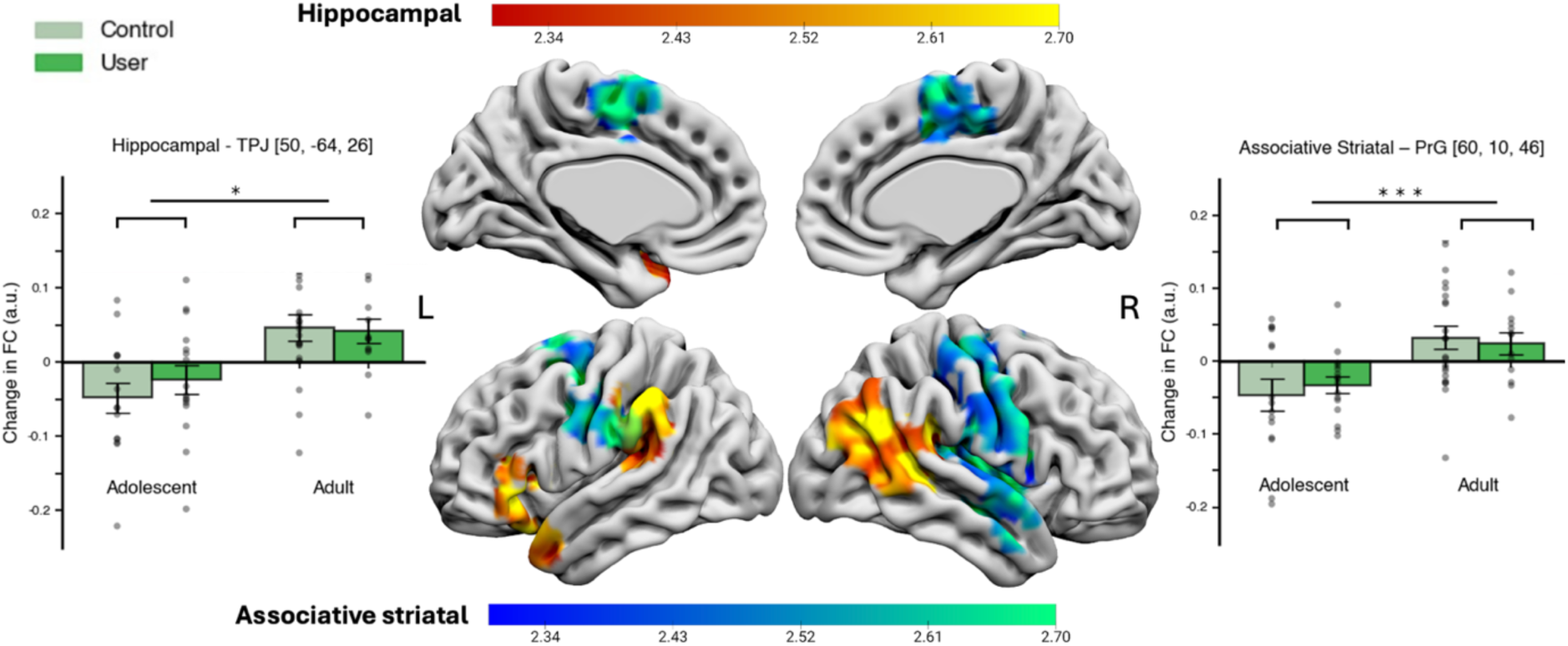
Main effects of age-group on the change in functional connectivity over one year in a seed-to-voxel resting-state analysis. Renders display lateral and medial left and right hemispheres of SurfIce BrainMesh_ICBM152 with overlays of significant clusters (*Z* > 2.3, *p* < 0.05) where change in functional connectivity significantly increased in adults and decreased in adolescents. Adjacent plots show representative group divergences using the most significant cluster per network, labeled with peak MNI coordinates. Significance cluster-based family wise error corrected: * *p* < 0.05, ** *p* < 0.01, ** *p* < 0.001 Hippocampal network: Left bar plot and red-yellow colour map overlay. Network to cluster decreases in functional connectivity in adolescents occurred into the temporo-parietal junction (TPJ) [50, −64, 26]; *Z* = 4.18, *p* < 0.001; left ventrolateral PFC [−44, 18, 10]; *Z* = 3.36, *p* = 0.048; and left supramarginal gyrus (anticorrelation strengthening in adolescents) [−64, −32, 46]; *Z* = 3.81, *p* = 0.030. Associative striatal network: Right bar plot and blue-green colour map overlay. Network increases in functional connectivity in adults to right precentral gyrus (PrG) dorsolateral prefrontal cortex [60, 10, 46]; *Z* = 3.78, *p* < 0.001); left PrG [−40, −10, 34]; *Z* = 3.89, *p* = 0.016; and medial premotor cortex [−6, −8, 58]; *Z* = 3.86, *p* = 0.019.

In the associative striatal network analyses, clusters were identified in the right dlPFC extending to lateral precentral and premotor cortices, left precentral gyrus, and medial premotor cortex (Table 3).

No main effects of age group were identified in any of the other networks tested.

### User group main effects

User group differences were identified in ECN and associative striatal network analyses. Connectivity between the ECN seed and clusters located bilaterally in the superior parietal lobule (SPL) decreased over the study period in cannabis users relative to controls, who showed increased connectivity. The observed bilateral SPL clusters extended into the medial precuneus and postcentral sulcus (Table 3). Figure 3 shows the significant clusters and corresponding subgroup FC changes over the study period (bar plots for remaining significant clusters are in Supplementary Figures S2–5).

**Figure 3.**
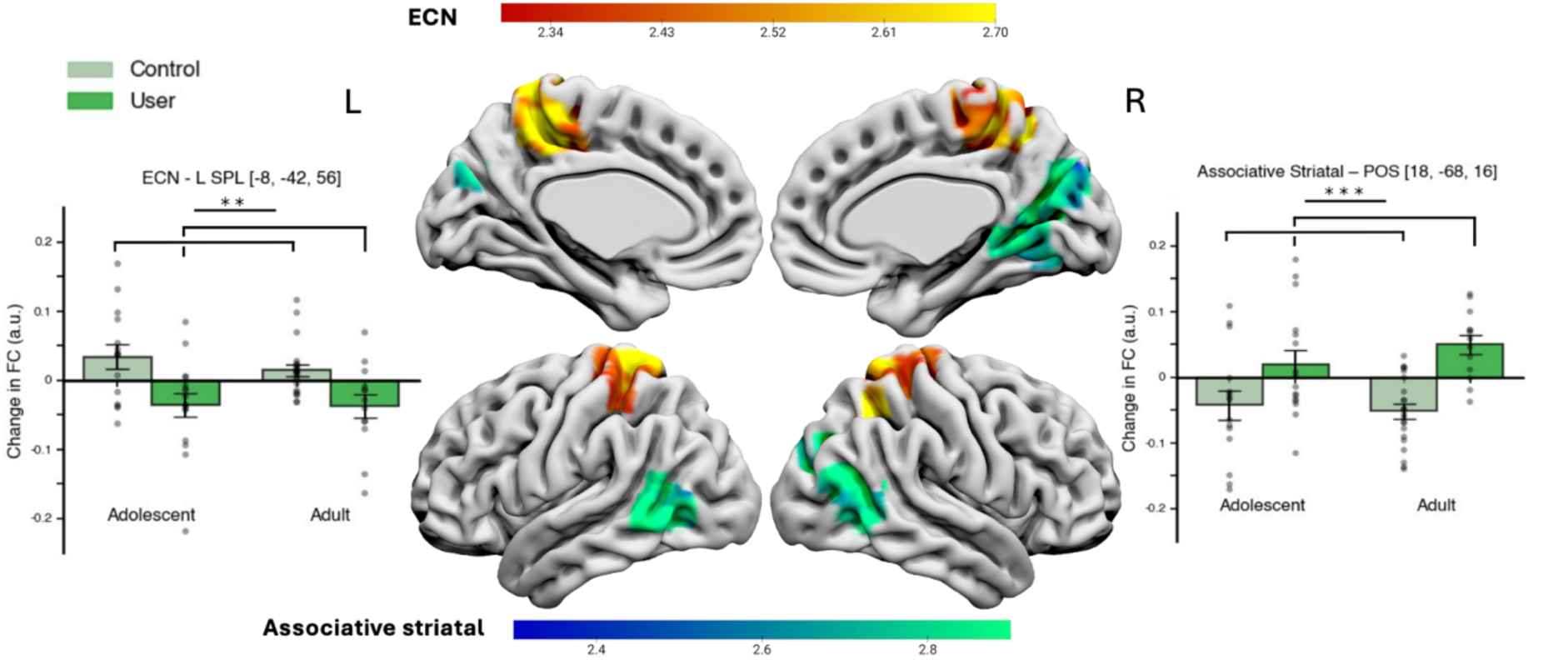
Main effects of user-group on the change in functional connectivity over one year in a seed-to-voxel resting-state analysis. Renders display medial and lateral left and right hemispheres of Surfice BrainMesh_ICBM152 with overlays of significant clusters (*Z* > 2.3, *p* < 0.05) where change in functional connectivity significantly differed between cannabis users and non-using controls. Adjacent plots show representative group divergences using the most significant cluster per network, labeled with peak MNI coordinates. Significance cluster-based family wise error corrected: * *p* < 0.05, ** *p* < 0.01, ** *p* < 0.001 Executive control network (ECN): Left bar plot and red-yellow colour map overlay. ECN decreased in functional connectivity with bilateral superior parietal lobule (SPL) in consistent cannabis users and increase in controls (L: [-8, –42, 56]; *Z* = 3.85, *p* = 0.004; R: [16, –54, 52]; *Z* = 3.69, *p* = 0.031). Associative striatal network: Right bar plot and blue-green colour map overlay. Network increased in functional connectivity (anticorrelation weakened) in consistent cannabis users and a decrease in functional connectivity (anticorrelation strengthened) in controls to the right parietal-occipital sulcus (POS) [18, −68, 16]; *Z* = 3.85, *p* < 0.001; and posterior MTG [-58, –58, 8]; *Z* = 3.59, *p* = 0.039.

FC changes that emerged between the associative striatal network and two temporo-occipital clusters were anticorrelated. Controls showed strengthened anticorrelation between these regions over time, whereas the anticorrelation weakened in cannabis users. The largest cluster was centred in the parieto-occipital sulcus, and a second cluster observed cluster was localised to the posterior middle temporal gyrus (MTG; Table 4).

No user group effects were identified in any of the other networks.

### Age × Cannabis Use Interaction Effects

Significant age x user group interactions were identified in the SN and DMN analyses (Table 4), with all significant clusters exhibiting negative connectivity with the seed. Interaction effects reflected divergent longitudinal trajectories between age and cannabis-use groups.

The majority of interaction effects showed strengthened anticorrelations in adult users and adolescent controls, with weakened anticorrelations in adult controls and adolescent users. This pattern was observed in two SN clusters, localised to the left dmPFC and right vmPFC, and one DMN cluster in the pre-supplementary motor area.

A second, opposing interaction pattern was observed in one SN cluster in the posterior mediodorsal thalamus and one DMN cluster in the cerebellum (lobules V/VI). These clusters showed strengthened anticorrelations in adult controls and adolescent users, whereas anticorrelations weakened in adult users and adolescent controls. In the cerebellar cluster, connectivity shifted from approximately zero functional relationship to negative adult controls and adolescent users, resulting in the emergence of anticorrelation over the study period.

Opposingly, in this cluster, adult users FC reduced to zero. These clusters and corresponding subgroup FC changes are shown in Figure 4 (bar plots for remaining significant clusters are in Supplementary Figures S2–5).

**Figure 4.**
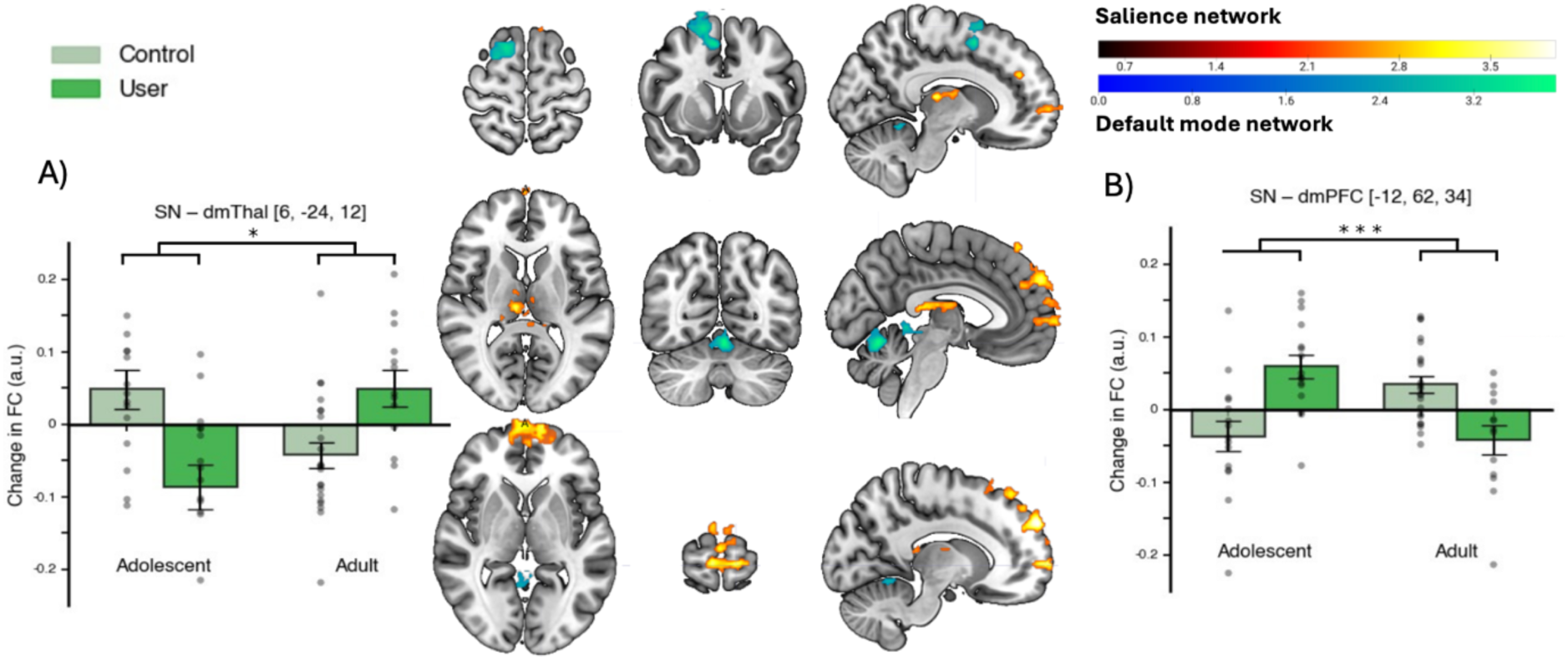
Interaction effects of age-group x user-group on the change in functional connectivity over one year in a seed-to-voxel resting-state analysis. Axial, coronal and sagittal slices of the MNI152 template in MRIcroGL overlain with significant clusters with bar plots showing divergent directional changes by group. All significant clusters were anticorrelated to their respective networks, an increase in functional connectivity represents a decrease in anticorrelation. Significance cluster-based family wise error corrected: * *p* < 0.05, ** *p* < 0.01, ** *p* < 0.001 Salience network: Clusters are overlaid in a red-to-yellow colour map. Right plot: dorsomedial prefrontal cortex (dmPFC) [ −12, 62, 34] and ventromedial PFC [−6, 42, 58] clusters strengthened in anticorrelation (respectively: *Z* = 4.13, *p* < 0.001; *Z* = 3.32, *p* = 0.042) in adult users and adolescent controls, with the opposite pattern by age and user group. Left plot: In a posterior mediodorsal thalamic cluster (dmThal) [6, −24, 12] adult controls and adolescent users showed strengthened anticorrelation at TP2, *Z* = 4.08, *p* = 0.024, with the opposite pattern in adult users and adolescent controls. Default mode network: Clusters are overlaid in a blue-to-green colour map. A cerebellar lobules V–VI [−4, −66, −14] cluster coupling to DMN mirrors directionality of plot A, *Z* = 3.94, p = 0.029. Pre-supplementary premotor area [6, 8, 52] showed the reverse pattern, *Z*= 3.74, *p* = 0.049, mirroring directionality in plot B. Bar plots for DMN interaction effects can be found in Supplementary Figure S5.

No interaction effects were identified in any other networks.

### Association with cannabis use

In the regression analyses, no significant associations were observed between the measures of cannabis exposure tested and longitudinal FC change in any significant cluster (Supplementary Figure S6).

## Discussion

The present findings suggest that regular cannabis use during adolescence is associated with altered developmental trajectories of large-scale functional network organisation. By directly comparing adolescent and adult users with their age-matched controls within a one-year longitudinal framework, these findings extend a predominantly cross-sectional cannabis neuroimaging literature by demonstrating that alterations differ according to developmental stage. Age-related changes evident in hippocampal and associative striatal networks were broadly consistent with ongoing neurodevelopment, however, cannabis-related effects converged on higher-order association systems. Specifically, users exhibited divergent longitudinal trajectories of salience and DMN connectivity with medial prefrontal and task-positive regions, alongside differences within executive and associative cortico-striatal circuitry when compared to their age-matched controls. Taken together, these findings indicate developmentally atypical changes of higher-order functional network architecture over a year occurring with regular cannabis usage and suggests that alterations preferentially involve distributed association systems rather than isolated regional abnormalities.

Among the age × user group interaction effects (Figure 4), SN effects were illustrative of altered functional coupling between salience-related and higher-order association regions, suggesting altered maturation of large-scale association systems. During normative adolescent development the SN is characterised by non-linear increases in functional specialisation and salience-mediated between-network integration, such as DMN and ECN switching (Gracia-Tabuenca et al., 2021; Lopez et al., 2020; Marek et al., 2015; Menon & Uddin, 2010). Against this backdrop, adolescent controls demonstrated increasing SN– mPFC anticorrelation over time, whereas adult controls showed reduced anticorrelation, potentially reflecting greater within-network specialisation in adulthood (Gracia-Tabuenca et al., 2021; Marek et al., 2015; Rubia, 2013). Users exhibited the opposite pattern at both developmental stages, suggesting divergence from their respective age-specific control trajectories.

Across age x group interaction effects, regional changes were often localised to canonical DMN, attentional, and task-positive territories rather than solely core SN itself, indicating that cannabis-related differences preferentially affect functional relationships within large-scale association systems. In adolescent controls, increasing DMN–task-positive anticorrelation is consistent with the strengthening of network segregation expected during late adolescent maturation (Marek et al., 2015; Rubia, 2013). The opposing trajectory observed in adolescent users, therefore, represents a departure from this age-matched developmental pattern. The strongest convergence with the present findings comes from Pujol et al. (2014) who reported enhanced default–insula anticorrelation in cannabis users, together with repeated observations of altered medial prefrontal, insular (/SN), and DMN connectivity (Filbey et al., 2018; Hammond et al., 2022; Wetherill et al., 2015; Wijayendran et al., 2018; Zhou et al., 2018). These findings might indicate a disruption of interactions among salience, default-mode, and executive systems.

Given the role of the mediodorsal thalamus in supporting cognitive control, reward-related processing, and large-scale network integration (Krebs et al., 2012; Seeley et al., 2007), the observed divergence in SN–thalamic functional connectivity further supports the interpretation of altered salience-related thalamocortical integration. Adolescent users increasing SN–thalamic anticorrelation in the opposite direction to mPFC coupling changes, a pattern consistent with a hypothetical progressive dissociation between thalamic relay function and downstream cortical control. Together, these convergent findings suggest that cannabis-related alterations preferentially involve distributed association and attentional network architecture rather than isolated abnormalities within individual functional systems.

Cannabis-related longitudinal alterations (main effects of user group; Figure 3) additionally converged on distributed executive-attentional systems, extending the evidence for altered association-network organisation beyond age-dependent trajectories observed for SN and DMN. This convergence suggests that the principal commonality across analyses lies not in the statistical pattern of effects, but in the neuroanatomical systems implicated, reinforcing that cannabis-related alterations localise to distributed association networks. One possible explanation for the absence of age × user group interactions in these networks is that prefrontal executive-attentional systems undergo protracted and heterogeneous pubertal-dependent maturation (Gracia-Tabuenca et al., 2021; Lopez et al., 2020; Marek et al., 2015; Rubia, 2013) potentially reducing a developmental dissociation between the age groups sampled here.

Within the ECN, cannabis users failed to demonstrate the strengthening of frontoparietal connectivity observed in controls, whereas associative corticostriatal analyses revealed attenuated development of anticorrelations with posterior transmodal cortices. Although anatomically distinct, these regions participate in distributed executive-attentional processing, supporting cognitive control, attentional allocation and flexible behavioural selection. The SPL and posterior MTG both occupy transmodal association cortex implicated in executive control, attentional allocation and semantic-action processing (Dixon et al., 2018; Seeley et al., 2007). Moreover, attentional-occipital effects align with prior reports of altered visual-attentional circuitry in cannabis users (Harding et al., 2012; Liu et al., 2020; Zhou et al., 2018). Although similar peripheral localisation has been interpreted as compensatory migration of network organisation (Ertl et al., 2023), the present findings favour an alternative interpretation whereby cannabis-related alterations emerge within transmodal regions that participate in salience attribution and multiple higher-order systems. One proposed mechanism of THC-induced psychosis is that endocannabinoid-mediated modulation of dopaminergic signalling in prefrontal and striatal circuits, may disrupt attentional, interoceptive, and higher-order cognitive processes, thereby altering reward processing, reinforcement learning, and salience attribution in ways that contribute to positive psychotic symptoms (Bloomfield et al., 2014; Fernandez-Espejo et al., 2009; Müller-Vahl & Emrich, 2008; Wijayendran et al., 2018). Collectively, these findings indicate that cannabis-related alterations extend beyond salience-mediated developmental interactions to encompass distributed executive-attentional circuitry, supporting a model in which cannabis disrupts strengthening of executive-attentional systems and the normative specialisation across higher-order association systems (Batalla et al., 2013; Blest-Hopley et al., 2018; Ertl et al., 2023; Hammond et al., 2022; Harding et al., 2012).

Several methodological considerations strengthen interpretation of the present findings. The observation of anticorrelational effects in the absence of global signal regression supports the biological validity of the reported negative coupling patterns (Chai et al., 2012). The longitudinal repeated-measures design reduced the inter-individual variability inherent to cross-sectional studies. Importantly, many cannabis-related effects were localised outside canonical network cores and instead involved transmodal regions participating in multiple large-scale systems. These distributed network patterns are difficult to capture using task-based paradigms, which target discrete functional mechanisms. Furthermore, the present findings reinforce the possibility that previous studies combining participants across periods of differing neurodevelopmental vulnerability may have obscured biologically meaningful differences in cannabis-related functional connectivity.

Several limitations should be considered when interpreting these findings. First, the relatively small sample size limited statistical power and constrained the extent to which potential confounding variables could be incorporated into the models. The limited sample size also restricted the ability to robustly evaluate dose–response relationships. Although urinary THC measures provided objective confirmation of recent cannabis exposure, frequency and quantity of cannabis consumption were characterised using self-report and therefore remain subject to recall or response bias. Moreover, as an observational study, pre-existing neurobiological differences between groups cannot be excluded and causal inferences regarding cannabis exposure are therefore limited. Second, the one-year follow-up provides a short window for developmental investigations. Although this interval captured measurable within-subject change, extended longitudinal follow-up will be necessary to more fully characterise the observed trajectory differences and any further functional or clinical consequences. Third, cannabis-related effects could not be fully dissociated from concomitant tobacco use, which independently affects rs-FC (Wetherill et al., 2015). While cannabis and tobacco are frequently co-used in naturalistic settings (Hindocha et al., 2016), this limits attribution of the observed effects to cannabis alone. Finally, the absence of robust ventral striatal findings should not be interpreted as supporting or opposing literature that suggests cannabis influences reward circuitry (Ernst et al., 2005; Lawn et al., 2016; van Hell et al., 2010; Yanes et al., 2018). The ventral striatum and adjacent subcortical regions are particularly vulnerable to susceptibility artefacts and physiological noise during fMRI acquisition (Srirangarajan et al., 2021), potentially reducing sensitivity to detect corticostriatal FC changes.

## Conclusion

The present findings demonstrate that cannabis-related alterations in longitudinal functional connectivity are developmentally dissociable and preferentially converge on distributed higher-order association systems. Adolescent users showed divergence from normative network maturation trajectories, whereas adult users demonstrated patterns consistent with reduced network segregation. Taken together our findings argue against a uniform effect of cannabis on brain organisation and supports a neurodevelopmentally sensitive framework. The adolescent window may represent a period of vulnerability, during which cannabis exposure perturbs the active refinement of higher-order association network refinement rather than disrupting an already-established architecture.

These findings carry broader clinical and translational relevance. Disruption of large-scale association network organisation, particularly involving salience-mediated interactions, is increasingly recognised as a transdiagnostic feature of neuropsychiatric disorders (Menon, 2011). Younger age of cannabis onset is among the strongest predictors of later psychosis risk (Arseneault et al., 2004), and the present findings offer a neurodevelopmental framework through which this epidemiological association may be understood. Specifically adolescent cannabis use may alter the developmental refinement of salience-mediated network organisation during a window where such perturbations carry long-term consequences. Future longitudinal work with an extended follow-up will be necessary to determine whether the trajectory divergences observed here are transient or persist into adulthood, and whether they are associated with functional and clinical sequelae.

## Data availability

Participants in the CannTeen study did not give consent to the sharing of their data outside the research team, and the data are therefore not available.

