## Supplementary Material for "Longitudinal fMRI reveals altered developmental trajectories of functional network organisation associated with adolescent cannabis use"

### Supplementary Methods

#### Note 1: MRI acquisition parameters

##### Structural

T1-weighted structural images were acquired using a magnetisation-prepared rapid gradient-echo sequence. TR = 2300 ms, TE = 2.98 ms, flip angle = 9°, 1 mm isotropic voxels.

##### Functional

T2*-weighted functional images were acquired using a multiband gradient-echo echo-planar imaging sequence. TR = 1250 ms, TE = 30 ms, flip angle = 62°, multiband acceleration factor = 2, GRAPPA = 2) with 44 interleaved slices and 3 × 3 × 3 mm isotropic voxels. Field of view was 192 mm, with an in-plane matrix of 64 × 64 voxels and a total of 384 volumes.

#### Note 2: MRI preprocessing

##### Structural workflow

T1-weighted images were preprocessed in FSL (Smith et al., 2004; Woolrich et al., 2001), including brain extraction (BET), bias field correction, tissue segmentation, and normalisation. Registrations for MRI data used FLIRT for linear registration, and FNIRT for nonlinear normalisation to MNI152 standard space (Andersson et al., 2007; Jenkinson et al., 2002). Tissue segmentation produced grey matter, white matter (WM), and cerebrospinal fluid (CSF) partial volume maps for each participant. Magnitude images were skull-stripped and combined with phase-difference images to generate voxel-wise fieldmaps.

##### Functional workflow

Functional images were pre-processed similarly to Ertl et al., (2023b) in FEAT using the following steps: head motion correction (MCFLIRT), temporal high-pass filtering (cutoff 100 s), spatial smoothing (6 mm FWHM Gaussian kernel), $B_0$ field unwarping using the brain-extracted fieldmaps. Registration to structural and standard space was performed using boundary-based registration and nonlinear warping to MNI152 2 mm space. A temporal low-pass filter (≈ 0.08 Hz implemented using $\sigma = 0.556$ with fslmaths) was then applied to remove high-frequency noise.

#### Seed regions for 7 networks of interest in MNI space

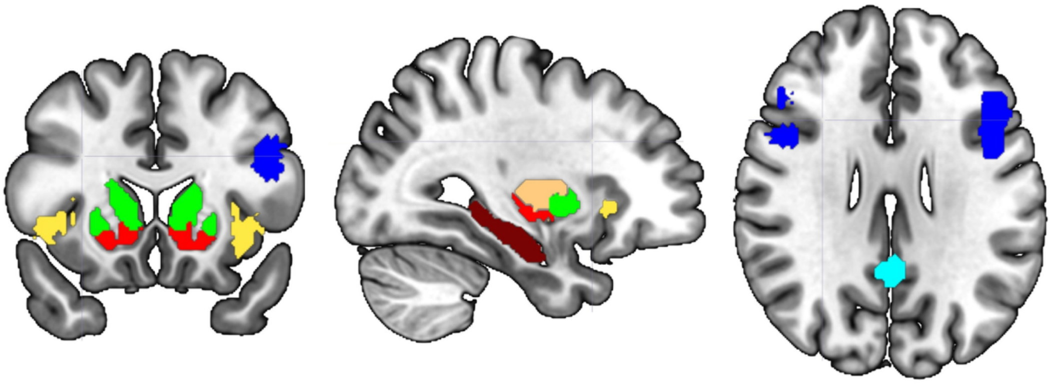

Figure S1. Coronal, sagittal, and axial slices of the MNI152 template overlain with the 7 different seed masks used for seed to voxel analysis of resting-state networks: hippocampus (brown), limbic striatum (ventral striatum; red), sensorimotor striatum (putamen; orange), salience (anterior insula; yellow), associative striatum (green), default mode (posterior cingulate cortex; cyan), executive control (dorsolateral prefrontal cortex; royal blue).

#### Supplementary Data Characteristics

#### Note 3: Participant Characteristics

#### Participant Demographics

Groups did not differ on sex, maternal education, own education, or ethnicity (all $p > 0.05$; see Table S2 for statistics).

##### *Sex and education*

Table S2. Demographic comparisons by cannabis user group and age group.

| | Controls | Users | Statistic | *p* |
|---|---|---|---|---|
| Gender, *n* (F/M) | 17/20 | 13/16 | $\chi^2 = 0.01$ | 0.928 |
| Mother's education, median | 4 | 6 | $U = 407$ | 0.135 |
| Own education, median | 5 | 3 | $U = 388$ | 0.074 |
| | **Adolescents** | **Adults** | **Statistic** | ***p*** |
| Gender, *n* (F/M) | 13/18 | 17/18 | $\chi^2 = 1.52$ | 0.218 |
| Mother's education, median[a] | 6 | 4 | $U = 390$ | 0.072 |

*Note*: [a]One adolescent excluded from mother's education analysis (data not available; remaining data *df* = 66). Own education not compared across age groups due to structural confound (adolescents cannot attain equivalent educational levels). Education was ranked on an ordinal 1–8 scale, 8 = highest.

##### *Ethnicity*

Table S3. Ethnic composition of cannabis user group and age group, with summary statistic of the percentage of non-White participants per group.

| | Ethnicity | | | | | |
|---|---|---|---|---|---|---|
| | Asian | Black | Mixed | Other | White | % non-White |
| Adolescent User* | 0 | 1 | 1 | 1 | 12 | 20 |
| Adolescent Control | 2 | 1 | 2 | 0 | 10 | 33 |
| Adult User | 1 | 1 | 1 | 1 | 9 | 31 |
| Adult Control | 5 | 2 | 0 | 1 | 14 | 36 |

*Note*: *One participants ethnicity data was not available. White vs. non-White composition did not differ significantly between groups, $\chi^2(3, N = 65) = 0.93$, $p = 0.818$.

#### Substance use measures

Cannabis users compared to controls (age collapsed) also differed in AUDIT scores and quantity of cigarettes per month ($p < 0.05$), however prevalence of tobacco smoking, frequency of alcohol consumed across a month, and BDI, BAI, and PSI scores at baseline did not differ by age or user group ($p > 0.05$).

##### *Tobacco*

##### User group prevalence

Table S4. Population prevalence of daily tobacco usage by user group.

| | Control | User | |
|---|---|---|---|
| Non-daily smokers/non-smokers | 34 | 23 | 57 |
| Daily smoker | 3 | 6 | 9 |
| Total n | 37 | 29 | 66 |

*Note*: Groups differed significantly in tobacco smoking prevalence ($\chi^2(66) = 2.19$, $p$ = 0.139), with a higher proportion of smokers among cannabis users than controls.

#### Age group prevalence

Table S5. Population prevalence of daily tobacco usage by user group.

| | Adolescent | Adult | |
|---|---|---|---|
| Non-daily smokers/non-smokers | 25 | 32 | 57 |
| Daily smoker | 6 | 3 | 9 |
| Total n | 31 | 35 | 66 |

*Note*: Tobacco smoking prevalence did not differ significantly between adolescents and adults ($\chi^2(66) = 1.62$, $p = 0.203$).

#### Cigarette use descriptive statistics

Table S6. Descriptive statistics of cigarette use between user groups (M, SD; and independent samples *t*-test).

| | Controls | Users | *df* | *t* | *p* |
|---|---|---|---|---|---|
| Cigarettes per month | 5 (10) | 15 (13) | 63 | 3.04 | 0.003 |
| Age of first use in years | 17.7 | 15.7 | 58 | 29 | 0.220 |

*Note*: Statistics were based on the data available. Cigarette use was higher in users than controls, but age of first cigarette use did not differ.

#### *Alcohol*

#### Days per month

Table S7. Descriptive statistics and independent samples *t*-test comparing age groups on alcohol use frequency.

| | Adolescents | Adults | *t* | *p* |
|---|---|---|---|---|
| Alcohol days per month | 3.5 (3.9) | 5.7 (6.1) | 1.72 | 0.090 |
| | Controls | Users | *t* | *p* |
| Alcohol days per month | 3.9 (4.22) | 5.59 (6.26) | 1.29 | 0.203 |

*Note*: Frequency given as an estimated mean number of days per month on which alcohol was consumed. *df* = 27

#### AUDIT

Table S8. Alcohol Use Disorders Identification Test (AUDIT) group descriptive statistics (M; SD) and independent samples *t*-test by user group.

| | Controls | Users | *df* | *t* | *p* |
|---|---|---|---|---|---|
| AUDIT | 4.1 (2.4) | 6.0 (3.4) | 57 | 2.50 | 0.015 |

*Note*: Statistics were based on the data available. Alcohol consumption was below hazardous thresholds across both groups but was slightly higher in the user group.

#### Mood and psychological scales

##### *Baseline Characteristics*

Repeated measures ANOVA for main effect of user group or age group (to preserve statistical power), showed no significant effects of BDI, BAI, or PSI of the year.

Table S9. Psychological scales baseline characteristics and mixed ANOVA for age and user group effects.

| | Adolescents | | Adults | | *Age effects* | | *User effects* | |
|---|---|---|---|---|---|---|---|---|
| | Controls | Users | Controls | Users | *F* | *p* | *F* | *p* |
| BDI | 11.6 (6.5) | 8.3 (5.3) | 7.1 (6.2) | 7.5 (8.8) | 2.59 | 0.112 | 0.77 | 0.384 |
| BAI | 12.9 (9.3) | 11.4 (8.5) | 7.6 (5.6) | 8.7 (9.4) | 3.89 | 0.053 | 0.009 | 0.923 |
| PSI | 27.9 (14.8) | 29.4 (19.7) | 19.8 (19.5) | 22.2 (15.4) | 2.94 | 0.091 | 0.178 | 0.674 |

Note: No interactions between age and user group were significant, *df* = 62. BDI = Becks Depression Inventory, BAI = Becks Anxiety Inventory, PSI = Psychotomimetic States Inventory.

#### Note 4: Data Quality Control

Attrition did not differ significantly by age group ($\chi^2$(1,66)= 0.58, $p$ = 0.448) or cannabis group ($\chi^2$(1,66) = 1.58, $p$ = 0.209). Mean framewise displacement (head movement) did not differ by age group ($F$(62)= 2.44, $p$ = 0.123) or cannabis group ($F$(62)= 0.92, $p$ = 0.340).

#### Supplementary Analysis Outcomes

#### Note 5: Complete results table (seed-to-voxel analysis)

Table S10. Summary of results across the 7 seed to voxel resting state analyses on year change data with significant and null effects of age group, user group, and age x user group interaction.

| Network | Effect | Location | Functional relationship | Direction |
|---|---|---|---|---|
| Salience | Age | – | – | – |
| | User | – | – | – |
| | Age × User group | Dorsomedial prefrontal cortex | Anticorrelation | AC+TU ↑; AU+TC ↓ |
| | | Ventromedial prefrontal cortex | Anticorrelation | AC+TU ↑; AU+TC ↓ |
| | | Dorsomedial thalamus | Anticorrelation | AU+TC ↑; TU+AC ↓ |
| Default Mode | Age | – | – | – |
| | User | – | – | – |
| | Age × User group | Cerebellar lobule V–VI | Anticorrelation | AU+TC ↑; TU+AC ↓ |
| | | Pre-supplementary motor area | Anticorrelation | AC+TU ↑; AU+TC ↓ |
| Executive Control | Age | – | – | – |
| | User | R Superior parietal lobule | Positive FC | C ↑, U ↓ |
| | | L Superior parietal lobule | Positive FC | C ↑, U ↓ |
| | Age × User group | – | – | – |
| Hippocampal | Age | Tempero-parietal junction | Positive FC | A ↑, T ↓ |
| | | Ventrolateral prefrontal cortex | Positive FC | A ↑, T ↓ |
| | | Anterior supramarginal gyrus | Anticorrelation | A ↑, T ↓ |
| | User | – | – | None |
| | Age × User group | – | – | None |
| Associative Striatal | Age | Dorsolateral prefrontal cortex | Positive FC | A ↑, T ↓ |
| | | Precentral gyrus | Positive FC | A ↑, T ↓ |
| | | Medial premotor cortex | Positive FC | A ↑, T ↓ |
| | User | Parieto-occipital sulcus | Positive FC | C ↓, U ↑ |
| | | Middle temporal gyrus (temporooccipital) | Positive FC | C ↓, U ↑ |
| | Age × User group | – | – | – |
| Limbic Striatal | Age | – | – | – |
| | User | – | – | – |
| | Age × User group | – | – | – |
| Sensorimotor Striatal | Age | – | – | – |
| | User | – | – | – |
| | Age × User group | – | – | – |

*Note*: Direction of change: A = Adults, T = Teenagers(/Adolescents), C = Controls, U = Cannabis users; * ↑ = increasing connectivity / reduced anticorrelation, ↓ = decreasing connectivity / strengthened anticorrelation over the year.

#### Note 6: Additional bar plots of functional change significant effects

##### Main effects of user group

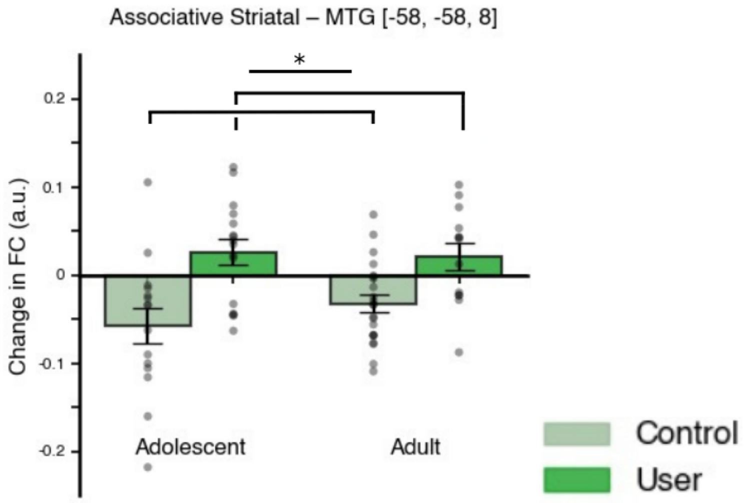

Figure S2. Bar plot of longitudinal resting state associative striatal network change in coupling to the posterior MTG [-58, -58, 8]. Network increased in functional connectivity (anticorrelation weakened) in consistent cannabis users and a decrease in functional connectivity (anticorrelation strengthened) in controls; $Z = 3.59$, $p = 0.039$. Significance cluster-based family wise error corrected: * $p < 0.05$, ** $p < 0.01$, ** $p < 0.001$.

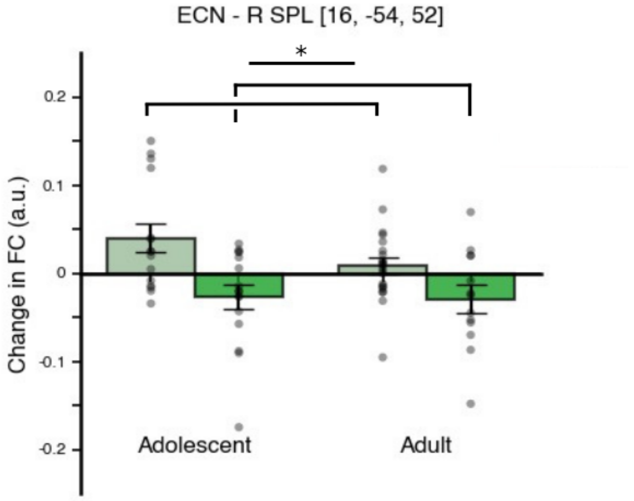

Figure S3. Bar plot of longitudinal resting state associative striatal network change in coupling with the right superior parietal lobule [16, -54, 52]. ECN decreased in functional connectivity within consistent cannabis users and increased in controls, $Z = 3.69$, $p = 0.031$. * $p < 0.05$, ** $p < 0.01$, ** $p < 0.001$.

#### Main effects of age group

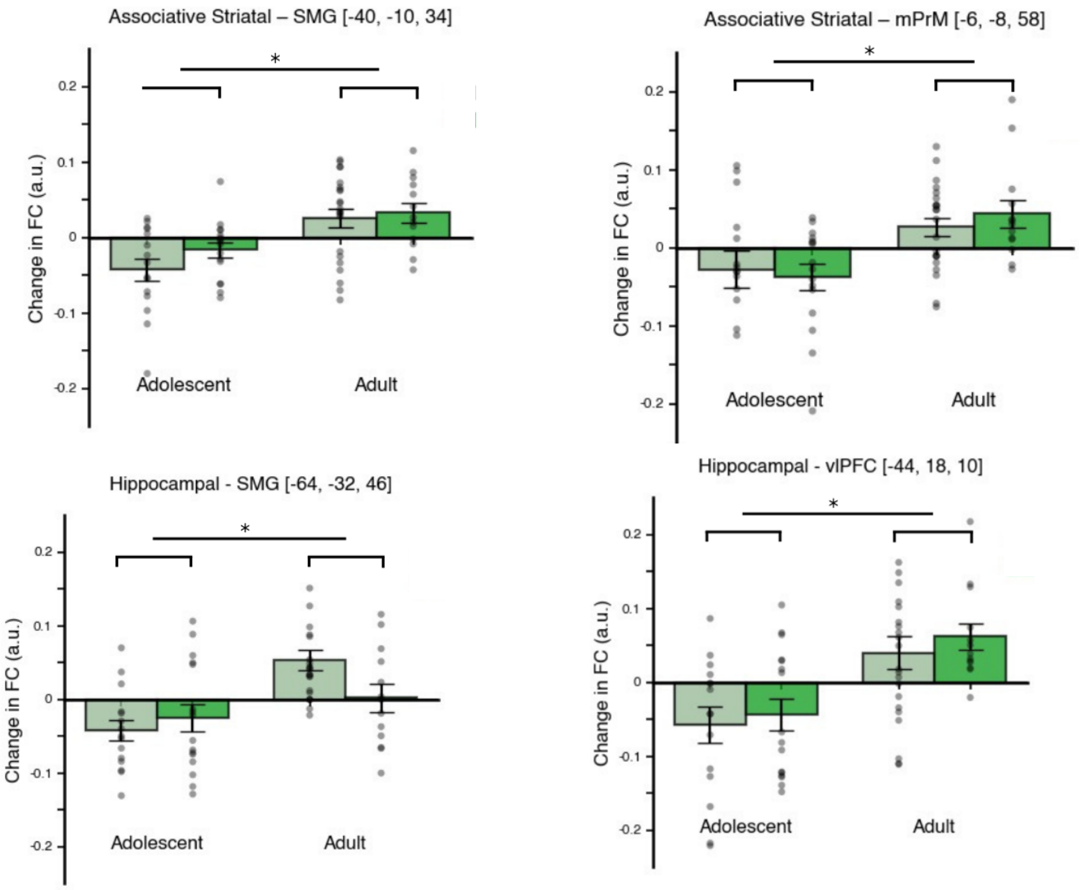

Figure S4. Bar plots showing main effect of age in longitudinal functional connectivity to the associative striatal network. Divergent age trajectories were identified in associative striatal coupling the left precentral gyrus [−40, −10, 34]; $Z = 3.89$, $p = 0.016$, and medial premotor cortex (mPrM) [−6, −8, 58]; $Z = 3.86$, $p = 0.019$. Significance cluster-based family wise error corrected: * $p < 0.05$, ** $p < 0.01$, ** $p < 0.001$.

#### Interaction effects of age and user group

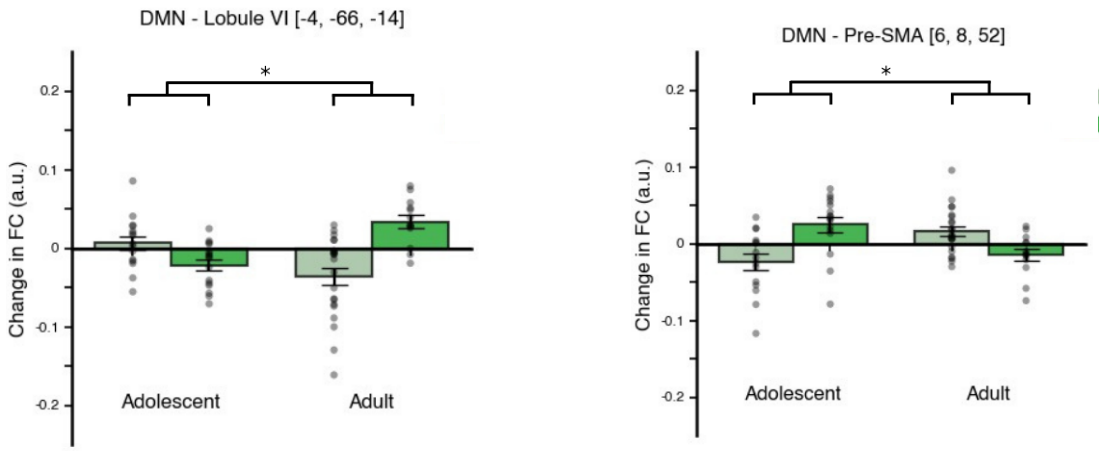

Figure S5. Bar plots showing divergent age by group changes in longitudinal functional connectivity were observed in DMN coupling. Left: cerebellar cluster lobules V/VI [−4, −66, −14] (anticorrelation weakening in adult users and adolescent controls, with the opposite effect in adult controls and adolescent cannabis users), $Z = 3.94$, $p = 0.029$. Right: pre-supplementary motor area (pre-SMA) [6, 8, 52] showed the reverse pattern, $Z = 3.74$, $p = 0.049$. Adolescent users and adult controls had weakened anticorrelations and adult users, and adolescent controls had strengthened to the DMN in this region. Significance cluster-based family wise error corrected: * $p < 0.05$, ** $p < 0.01$, ** $p < 0.001$.

#### Note 6: Functional connectivity change association with cannabis use frequency

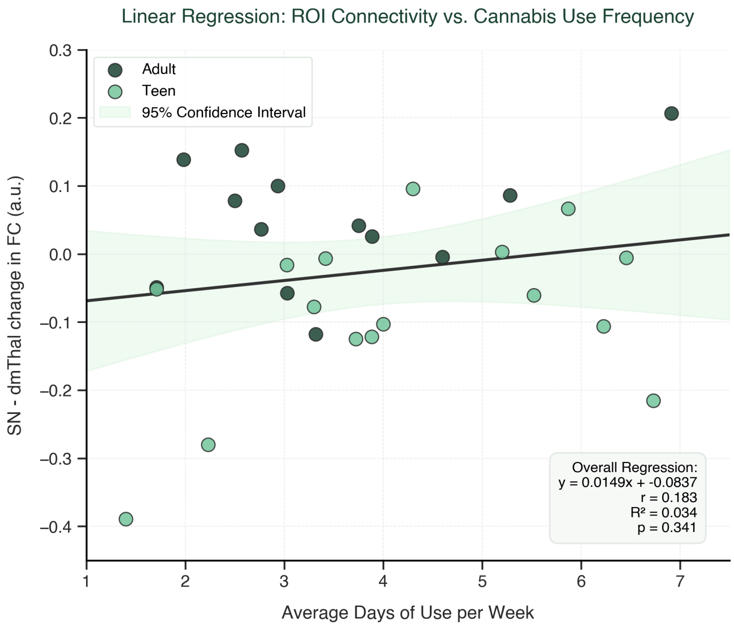

Figure S6. Representative scatter plot of significant FC change ROI value against average cannabis usage in days per week. Dorsomedial thalamus [6, 24, 12] cluster in the salience seeded network analysis change in functional connectivity in arbitrary units over the year in a linear regression against frequency of cannabis usage in days per week.
